## Supplemental Material for "Splicing Factor SRSF1 Deficiency in the Liver Triggers NASH-like Pathology via R-Loop Induced DNA Damage and Cell Death"

### Supplementary Methods

**eCLIP-seq library preparation, data processing, and peak calling.** Isolated hepatocytes suspended in ice-cold 1X PBS were crosslinked with 400 mJ/cm$^2$ of 254 nm UV radiation to stabilize RNA binding protein (RBP)–RNA interactions. Subsequent immunoprecipitation of SRSF1-RNA complexes, RNA isolation, library preparation and sequencing were performed as described previously (*84*). Briefly, crosslinked cells were lysed in buffer and sonicated, followed by treatment with RNase I (Thermo Fisher) to fragment RNA. SRSF1 antibody (A302-052A Bethyl Labs) were pre-coupled to anti-rabbit IgG Dynabeads (Thermo Fisher), added to lysate, and incubated overnight at 4 °C. Prior to IP washes, 2% of sample was removed to serve as the paired input sample. For IP samples, high- and low-salt washes were performed, after which RNA was dephosphorylated with FastAP (Thermo Fisher) and T4 PNK (NEB) at low pH, and a 3′ RNA adaptor was ligated with T4 RNA ligase (NEB). Ten per cent of IP and input samples were run on an analytical PAGE Bis-Tris protein gel, transferred to PVDF membrane, blocked in 5% dry milk in TBST, incubated with SRSF1 antibody used for IP (typically at 1:4,000 dilution), washed, incubated with HRP-conjugated anti-Rabbit secondary TrueBlot antibody (Rockland), and visualized with standard enhanced chemiluminescence imaging to validate successful IP. Ninety per cent of IP and input samples were run on an analytical PAGE Bis-Tris protein gel and transferred to nitrocellulose membranes, after which the region from the protein size to 75 kDa above protein size was excised from the membrane, treated with proteinase K (NEB) to release RNA, and concentrated by column purification (Zymo). Input samples were then dephosphorylated with FastAP (Thermo Fisher) and T4 PNK (NEB) at low pH, and a 3′ RNA adaptor was ligated with T4 RNA ligase (NEB) to synchronize with IP samples. Reverse transcription was then performed with AffinityScript (Agilent), followed by ExoSAP-IT (Affymetrix) treatment to remove unincorporated primer. RNA was then degraded by alkaline hydrolysis, and a 3′ DNA adaptor was ligated with T4 RNA ligase (NEB). qPCR was then used to determine the required amplification, followed by PCR with Q5 (NEB) and gel electrophoresis to size-select the final library. Libraries were sequenced on the HiSeq 4000 platform (Illumina). eCLIP was performed on IP from two independent hepatocyte samples from wildtype C57BL/6j mice, along with paired size-matched input before the IP washes. eCLIP data was processed using the rigorous eCLIP processing pipeline as described previously. A detailed description of steps and scripts used is available on the ENCODE website at https://www.encodeproject.org/documents/3b1b2762-269a-4978-902e-0e1f91615782/@@download/attachment/eCLIP_analysisSOP_v2.0.pdf.

Briefly, raw eCLIP reads were demultiplexed, adapter trimmed (cutadapt v1.9.dev1), and dropped if less than 18 bp in length. Processed reads where then mapped to the mm9 mouse genome using STAR (v2.4.0i) and filtered for duplicated reads. Peak calling was performed on usable reads considering only read 2 (read that is enriched for termination at crosslink site) using the publicly available tool CLIPper (available at https://github.com/YeoLab/clipper/releases/tag/1.0) with options –s mm9 –o –bonferroni –superlocal --threshold-method binomial --save-pickle (*42*). Reproducible and significantly enriched peaks were identified using a modified IDR method as described previously (*85*). Replicate-merged peaks with an IDR cutoff of 0.01 as well as $P \leq 0.001$ and fold enrichment ≥8 (using the geometric mean of $\log_2$(fold enrichment) between the two replicates) were considered reproducible and significant. All subsequent analysis was performed using the significant peak set.

**eCLIP peak annotation and analysis.** Peak annotation was performed by overlapping peak coordinates with feature coordinates in the mouse vM19 annotation available from GENCODE. Motif enrichment analysis within SRSF1 binding peak regions was performed using the

STREME function, found in The MEME Suite (v5.3.3, https://meme-suite.org/meme/index.html), using standard parameters (*86*). Sequences of the binding peak regions were obtained using the getFasta function available in bedtools (2.30.0) on the peak coordinates. Gene ontology analysis was performed using Enrichr on the genes with at least one associated transcript containing at least one SRSF1 binding peak. Overlap between SRSF1 binding peaks and DEGs was defined as at least one DEG associated transcript containing at least one SRSF1 binding peak. Overlap of SRSF1 binding peaks with differentially spliced exon was defined as the peak being within the region between the constitutive upstream and downstream exon relative to the alternative exon.

To determine if SRSF1 binding was resulting in significant differential splicing response, metanalysis of SRSF1 binding peak distribution was performed on differentially spliced cassette exons (SE events; n = 2,040) identified using rMATS in the acSRSF1 HKO model. The analysis approach was adapted from a previously described method (*87*). Peak signals were identified for 500 nt window flanking each exon boundary, extending a maximum of 100 nt into each exon and 400 nt into each intron. For shorter exons (<200 nt) and introns (<400 nt), signal was counted until the boundary of the neighboring feature. This creates two windows comprising of the 5' and 3' end of the cassette exon, resulting in a total vectorized region of 1000 nt. Each position within each vectorized region was marked 1 if it within a significant peak, and 0 otherwise. The values at each position were then summed and divided by the total number of events at each position to obtain the final peak density distribution. A peak distribution was also obtained for the native cassette exons (n = 2,013) which are exons found to be alternative (0.05 < PSI < 0.95) but do not change significantly ($|\Delta PSI| \leq 0.05$) in acSRSF1 HKO. Finally, significance and confidence intervals were determined based on a bootstrapping approach. Peak distribution was calculated for a random sample of *n* events (without replacement) from a set of background constitutive exons (n = 23,905; PSI > 0.95 and |ΔPSI| ≤ 0.05), where *n* is the number of significant SE events (n = 2,040). This was repeated 1,000 times to create a distribution of peak density at each nt position. The 95% confidence interval at each position was determined to be the range bounded by the $2.5^{th}$ and $97.5^{th}$ percentile values. This interval was used to identify the positions where RBP-responsive event maps were significantly different than native events.

**Analysis of ENCODE datasets for SR protein knockdowns.** Analysis of publicly available shRNA-seq datasets for SR proteins in HepG2 cells from the ENCODE project were performed in the same manner as described in *RNA-seq library preparation, sequencing, and analysis.* Briefly, differential gene expression analysis was performed using kallisto, tximport and DESeq2 packages on paired-end fastq files for each sample. Differential splicing analysis was performed using rMATS on available bam alignment files. Finally, gene ontology analysis was performed using gProfiler, a web-based gene ontology analysis tool. Details regarding the datasets used for this analysis are listed in *Supplementary Table 5*. Briefly, ENCODE datasets of SRSF1, SRSF3, SRSF5, SRSF7 and SRSF9 protein knockdown by shRNA in HepG2 cells were used for this study (*59*).

**Immunofluorescent staining assays in cultured cells**. For immunofluorescent staining, cells were fixed for 10 minutes using 4% PFA solution followed by a PBS wash three times and then permeabilized with 0.2% Triton X-100 plus 1% normal goat serum (NGS) in PBS/pH 7.3 for 5 minutes on ice. Cells were then washed with PBS with 1% NGS and then incubated in primary antibody dilution for 1 hour at room temperature in a humidified chamber. This was followed by another wash and then incubation with secondary antibody dilution for 1 hour at room temperature. The cells were then washed with PBS and stained with DAPI before inverting the coverslip onto a glass slide with aqueous mounting media. All antibodies used and respective dilutions are listed in Supplementary Table 3. For the fluorescent protein synthesis assay, the Click-iT™ HPG Alexa Fluor™ 488 Protein Synthesis Assay Kit (Thermo Scientific #C10428) was used following manufacturer's instructions. Briefly, 2 hours prior to harvesting, culture media was replaced with methionine-free DMEM media. Cells were incubated in media for 1 hour before supplementing well media with 50 μM of ClickIT-HPG reagent. Cells were incubated for additional 1 hour before fixation. Fixed cells were prepared following standard kit protocol. For the fluorescent Annexin V cell death staining, the Dead Cell Apoptosis Kit with Annexin V FITC and PI (Thermo Scientific #V13242) was used following manufactures instructions.

**Polysome gradient fractionation.** Isolated hepatocyte samples were prepared for polysome profiling following a previously described protocol (*88*). Briefly, hepatocytes were isolated as described previously, however, all buffers were supplemented with 150 μg/mL cycloheximide. The pellets were flash frozen in liquid nitrogen and stored at -80 °C until ready for polysome fractionation. Frozen hepatocytes were thawed on ice for 15 minutes with 1 mL of polysome lysis buffer containing 10 mM Tris-HCl (pH 8.0), 150 mM NaCl, 5 mM $MgCl_2$, 1% Nonidet-P40, 40 mM dithiothreitol, 1 U/mL SUPERaseIn RNase inhibitor (Thermo Fisher) and 150 μg/mL cycloheximide. Thawed cells were pipetted gently 10 times to ensure lysis of cytoplasm. The cell nuclei and debris were removed by centrifugation at 12,000 x g for 1 minute at 4 °C. The supernatant was transferred to a fresh tube and then centrifuged again at 16,000 x g for 7.5

minutes at 4 °C to remove remaining cell debris and organelles. The resulting supernatant was transferred to a fresh tube and about 400 µL supernatant was layered onto a 12 mL linear sucrose gradient (10 – 50% sucrose (w/v) made using a Biocomp Gradient Master) and centrifuged in an SW-41Ti rotor (Beckman) for 125 minutes at 38,000 r.p.m. at 4 °C. The fractionated sample was gently moved through the detector using a peristaltic pump set at 3.5 mL/min with 60% sucrose as the chase solution. Polysome profiles were measured with a UA-6 absorbance (ISCO) detector at 254 nm and recorded using the associated Peak Chart software.

**Serum fractionation for lipoprotein particle analysis.** Plasma was collected from mice after a 6 hour fast starting from noon and ending at 6 p.m. Plasma (~ 200 µL) was injected onto a Superose HR6 10/300 GL FPLC column (GE Healthcare). Lipoproteins were eluted with 24 mL of elution buffer containing 0.15 M NaCl, 1 mM EDTA, 0.2% w/v of sodium azide in PBS and 0.5 mL fractions were collected at a flow rate of 0.5 mL/min. Triglyceride and cholesterol concentration was measured using Infinity kits (Thermo Scientific). In a microtiter plate, 100 µL of plasma as well as standards were incubated with 100 µL of Infinity reagent and then incubated at 37 °C for 30 minutes. The plates were measured for absorbance using a BioTek machine at 500 nm. Using absorbance measurements, the quantity of triglyceride and cholesterol were calculated.

**Glucose tolerance test**. Mice were fasted overnight and injected with D-glucose at 2g/kg intraperitoneally. Blood glucose concentrations were measured at 0, 15, 30, 45, 60 and 120 min after glucose injections using the One Touch glucose meter. Blood was obtained from tail tip after clipping.

**Total RNA isolation and quantitative RT-PCR**. Total RNA was isolated from about 50 mg of snap frozen liver tissue or snap frozen cells from 6-well culture plates with TRIzol (Invitrogen) using the protocol described in the manual. Quality of the RNA was assessed by running about 1 µg of RNA on a bleach gel to evaluate the 28S and 18S bands (*89*). Approximately 5 µg of RNA was then reverse-transcribed into cDNA using the Maxima Reverse Transcriptase (Thermo Fisher Scientific) following manufactures protocol. Relative gene expression analysis was performed with ~50 ng of cDNA per reaction using a SYBR® Green™ based assay for Real-Time Quantitative Reverse Transcription PCR (qRT-PCR) using standard cycling conditions. Using *36B4* as a loading control, relative gene expression compared to the control group was calculated by the double Ct method (*90*). Primers sequences used for qPCR and splicing assays are listed in Supplementary Table S4.

**Xbp1 splice isoform analysis.** Total RNA and cDNA were prepared as specified as described in the section *Total RNA isolation and quantitative RT-PCR*. The cDNA was diluted to 25 ng $\mu l^{-1}$ with nuclease free water from which 1 µL would be used for PCR reactions. PCR reactions were setup using Taq DNA Polymerase (NEB #M0237) along with forward and reverse primers to a final concentration of 0.2 µM. Cycling conditions used for the PCR reaction are as follows; Initial denaturation at 95 °C for 30 seconds, 30 cycles consisting of denaturation at 95 °C for 20 seconds, annealing at 60 °C for 30 seconds, and extension at 68 °C for 30 seconds, and a final extension at 68 °C for 5 minutes. For visualization, 10 µL of reaction mixture was mixed with loading dye and then resolved on a 5% PAGE gel, stained in an ethidium bromide bath, and then imaged using ChemiDoc XRS+.

**TUNEL staining for apoptosis**. TUNEL staining was performed on paraffin tissue sections that were deparaffinized and rehydrated using standard procedure. Staining was performed with the *In-Situ Cell Death Detection Kit, Fluorescein* (Sigma Cat# 11684795910) using manufacturer protocol. Briefly, rehydrated tissue sections were permeabilized and then labelled using the TUNEL reaction mixture from the kit. The sections were incubated with the reaction mixture for 1 hour at 37 °C in a humidified chamber. Slides were then rinsed with PBS three times, counterstained for nucleus, and mounted with coverslip. For positive control, the section was treated with 20 U of DNAse (NEB) mix for 15 minutes. The slides were imaged using a confocal microscope.

### Supplementary Figures

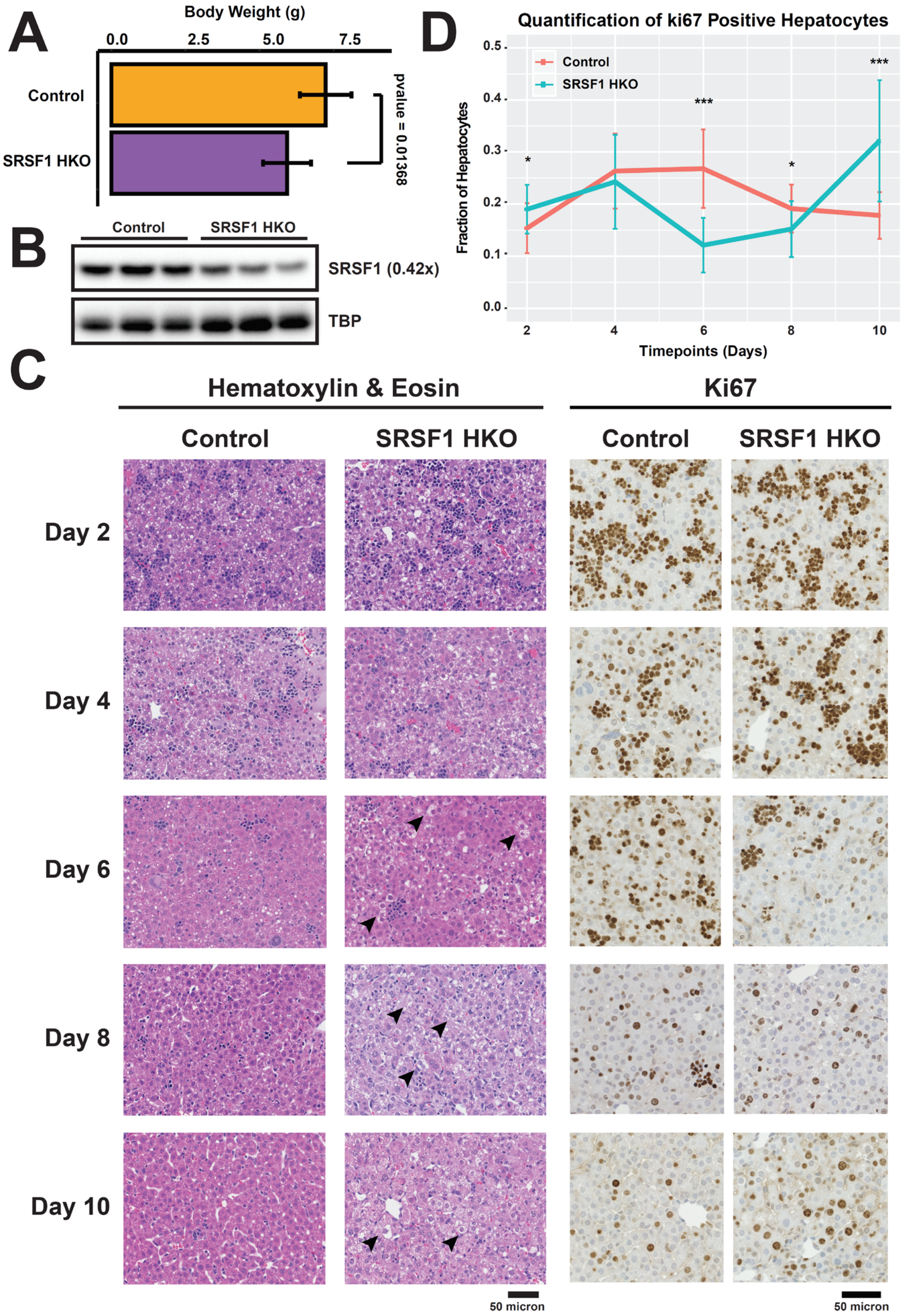


**Fig S1: Supplementary Figure S1 related to Figure 1.**

**Phenotypic characterization of SRSF1 HKO at post-natal timepoints.** (**A**) Total body weight measurements at post-natal day 10 (PN10) of SRSF1 HKO and Control mice (n = 6 per group). **(B)** Representative western blot image for relative abundance measurements of SRSF1 in hepatocytes isolated from PN10 SRSF1 HKO and Control mice with TBP as loading control. Value in parentheses signifies abundance fold change in SRSF1 HKO relative to Control. **(C)** Hematoxylin &

Eosin (*Left)* and Ki67 antigen, a proliferation marker, immunohistochemical *(Right)* staining of liver sections from SRSF1 HKO and Control mice at the indicated timepoint. Black arrows indicate ballooning necrotic hepatocytes. **(D)** Quantification of the fraction of Ki67-positive hepatocytes from staining images for each group and timepoint. Values are displayed as mean ± SD. *$P < 0.05$, **$P < 0.01$, ***$P < 0.001$.

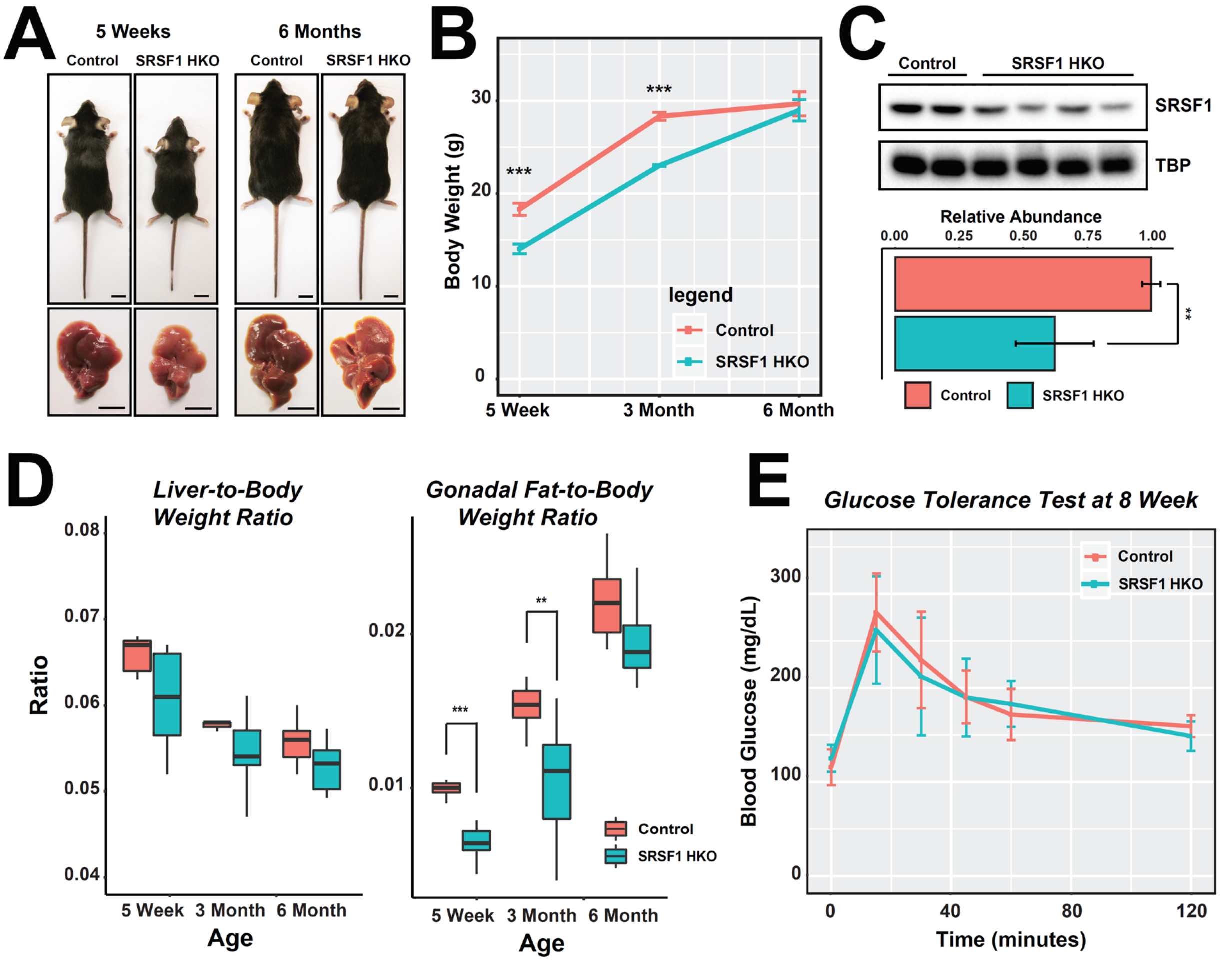


**Fig S2: Supplementary Figure S2 related to Figure 2.**

**Phenotypic differences in SRSF1 HKO mice progressively normalize to Control mice. (A)** Gross appearance of control and SRSF1 HKO mice and livers at 5 weeks and 6 months timepoints. Scale bar = 1 cm. **(B)** Body weight growth curve of controls and SRSF1 HKO mice. **(C)** Relative SRSF1 protein abundance quantification in SRSF1 HKO mice at 5 weeks using western blot analysis. **(D)** Box plot of liver-to-body weight and gonadal fat-to-body weight ratios in control and SRSF1 HKO mice at indicated timepoints during adult development. **(E)** Intraperitoneal glucose tolerance test of control and SRSF1 HKO mice at 8 weeks of age (n = 10 – 11 per group). Area under curve p-value = 0.759. Values are displayed as mean ± SD. Two-tailed unpaired *t* test was used to determine significance between 2 groups. *$P < 0.05$, **$P < 0.01$, ***$P < 0.001$.

| Serum Parameter | Age | Control | SRSF1 HKO | P-Value | Direction |
|---|---|---|---|---|---|
| Blood Glucose (mg/dL) | 1 Month | 118.7 ± 3.8 | 121.7 ± 6.8 | 0.2925 | N.S. |
| | 3 Month | 129.4 ± 4.6 | 118.7 ± 9.3 | 0.0242 | Down |
| | 6 Month | 138.7 ± 19.7 | 125.0 ± 20.6 | 0.2186 | N.S. |
| ALT (U/L) | 1 Month | 19.2 ± 8.5 | 231.1 ± 58.7 | 0.0001 | Up |
| | 3 Month | 13.9 ± 5.6 | 27.8 ± 12.1 | 0.0474 | Up |
| | 6 Month | 9.1 ± 5.2 | 15.4 ± 5.6 | 0.1005 | N.S. |
| AST (U/L) | 1 Month | 43.1 ± 31.2 | 225.2 ± 39.4 | 0.0001 | Up |
| | 3 Month | 25.7 ± 7.8 | 89.8 ± 17.0 | 0.0002 | Up |
| | 6 Month | 41.1 ± 10.5 | 42.5 ± 18.5 | 0.9018 | N.S. |
| Triglyceride (mg/dL) | 1 Month | 80.4 ± 13.7 | 65.5 ± 2.9 | 0.0277 | Down |
| | 3 Month | 143.7 ± 10.6 | 100.5 ± 12.6 | 0.0000 | Down |
| | 6 Month | 115.9 ± 10.7 | 128.9 ± 22.1 | 0.2020 | N.S. |
| Cholesterol (mg/dL) | 1 Month | 119.2 ± 22.7 | 90.2 ± 9.9 | 0.0343 | Down |
| | 3 Month | 96.7 ± 22.4 | 120.6 ± 17.3 | 0.0350 | Down |
| | 6 Month | 103.1 ± 8.0 | 121.5 ± 21.0 | 0.0612 | N.S. |

**Table S1: Supplementary Table S1 related to Figure 3.**

**Measurement of serum metabolic and liver function markers in SRSF1 HKO and Control mice at the indicated ages.** Mice were fasted for 10 hours prior to collection of serum (n = 7 per group)

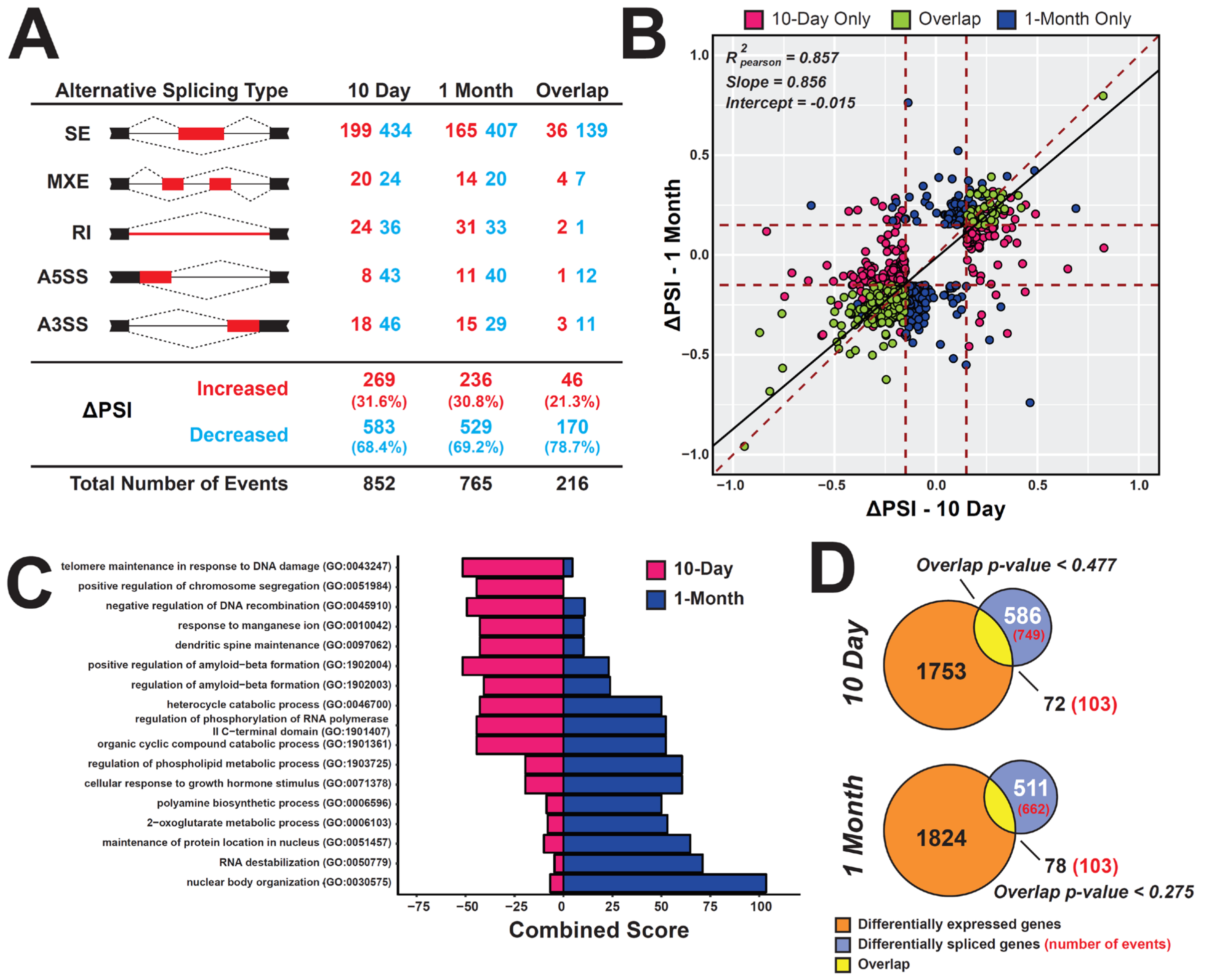


**Fig S3: Supplementary Figure S3 related to Figure 4.**

**RNA-seq analysis of SRSF1 HKO reveal extensive changes in exon usage. (A)** Breakdown of the number of differentially spliced exons (FDR < 0.10, junction read counts ≥ 10, and difference in Percent Spliced Index | ΔPSI| > 15%) by event type in 10-day and 1-month SRSF1 HKO mice. Numbers in red signify increased inclusion while blue signify decreased inclusion in SRSF1 HKO. SE, skipped exon; MXE, mutually exclusive exons; RI, retained intron; A5SS, alternative 5' splice site; A3SS, alternative 3' splice site. **(B)** Scatter plot showing the distribution of ΔPSI values for differentially spliced exons in 10-day and 1-month SRSF1 HKO mice. **(C)** Enrichr gene ontology analysis using genes with differentially spliced exons in 10-day and 1-month SRSF1 HKO. Larger combined score values signify greater enrichment. **(D)** Venn diagrams depicting overlap of genes with differential expression and splicing at each timepoint.

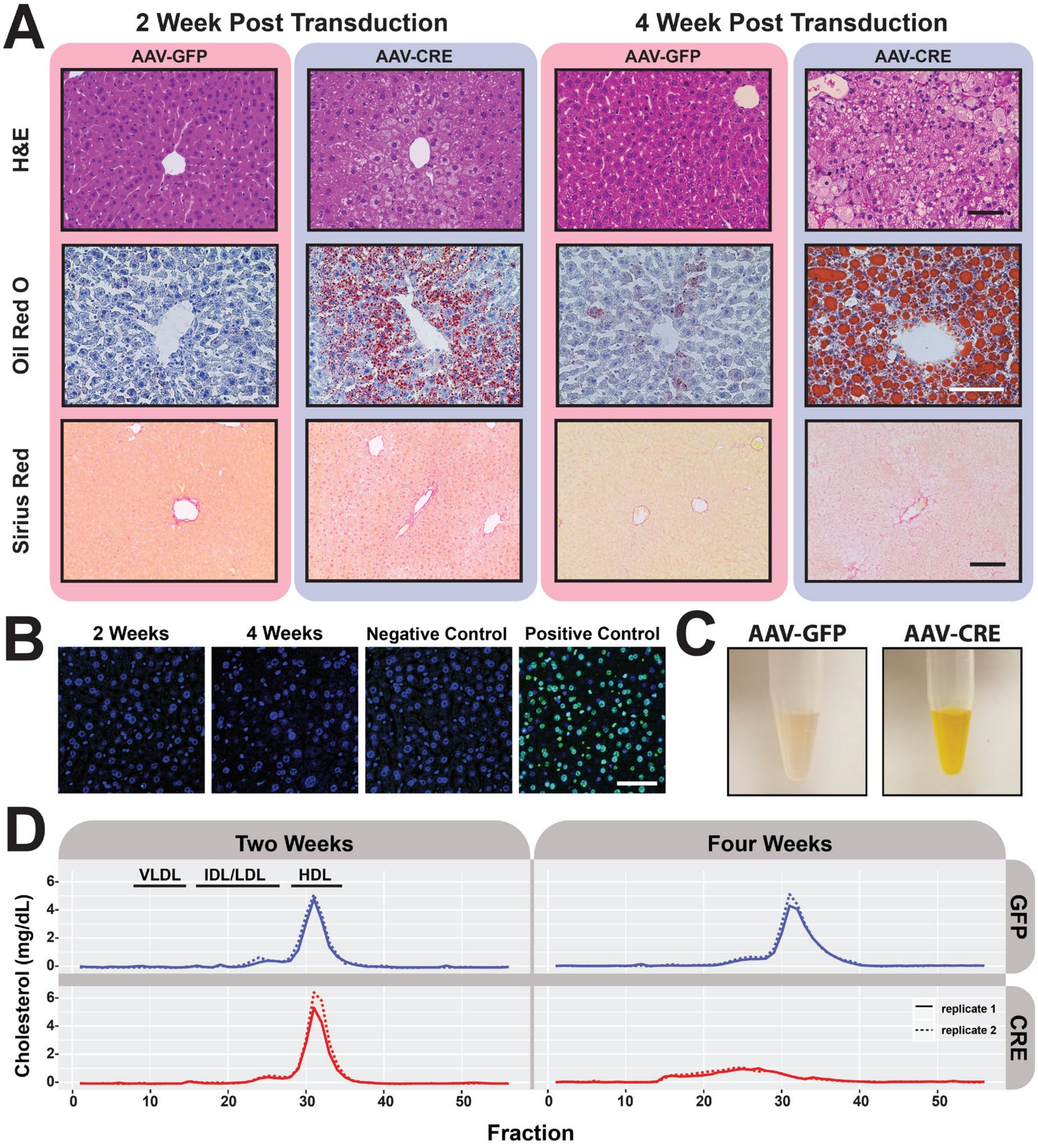


**Fig S4: Supplementary Figure S4 related to Figure 5 and Figure 6.**

**Acute loss of SRSF1 in adult hepatocytes recapitulate SRSF1 HKO phenotype. (A)** Representative images of H&E, Oil Red O (red, neutral lipids), and Sirius Red (red, collagen) histological staining of liver tissue harvested from control and acSRSF1 HKO mice at the indicated timepoints (n = 6 per group). Scale bar = 100 micron. **(B)** TUNEL staining for apoptosis detection in acSRSF1 HKO mice at 2 and 4 weeks. Green nuclei indicate apoptotic cells. **(C)** Representative images of serum collected from control (AAV-GFP) and acSRSF1 HKO (AAV-CRE) 4-weeks post viral transduction of SRSF1$^{flox/flox}$ mice. **(D)** Serum lipoprotein particle fractionation from 6 hour fasted control and acSRSF1 HKO mice. Each replicate consists of serum pooled from 2 biological replicates.

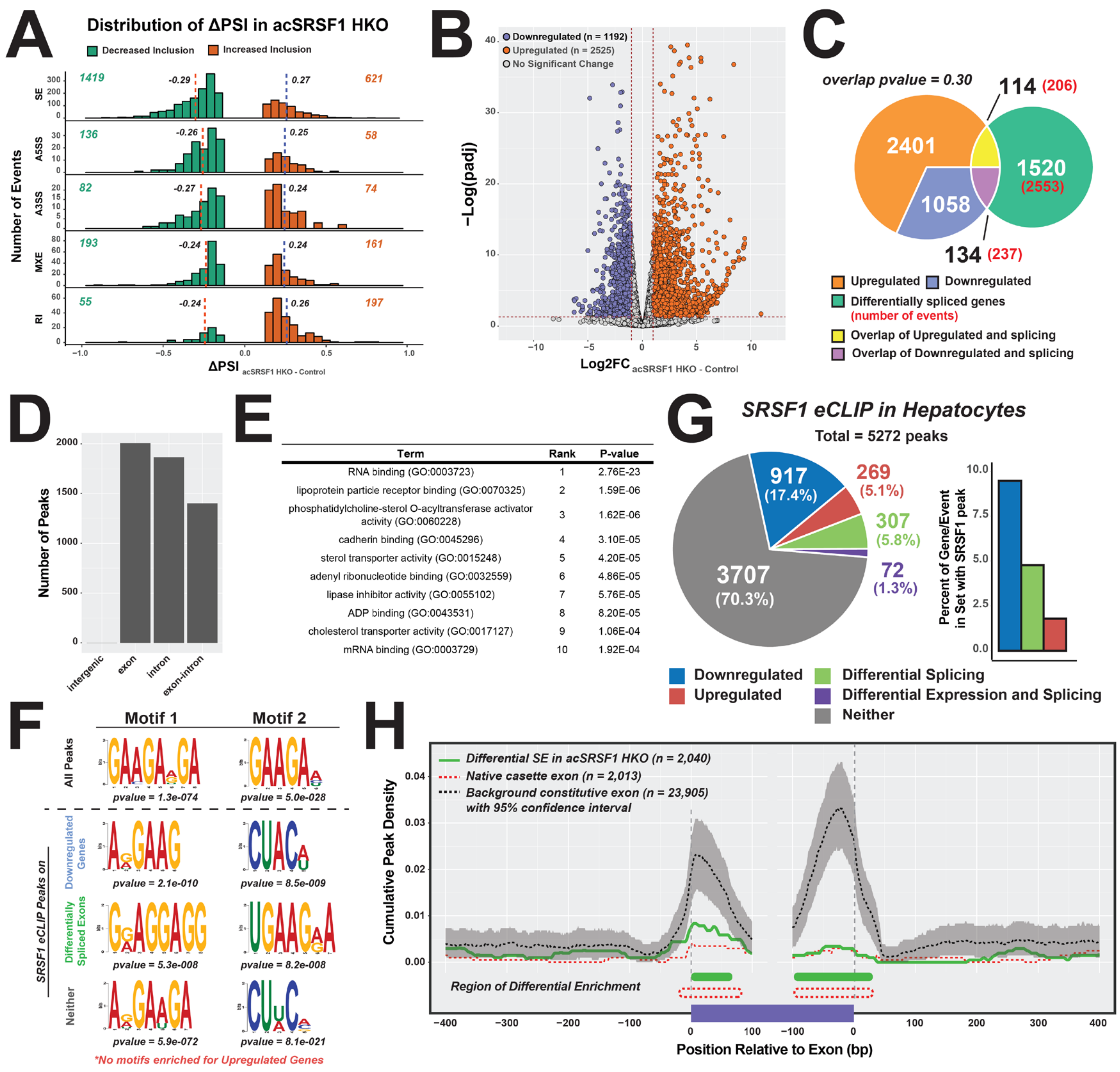


**Fig S5: Supplementary Figure S5 related to Figure 7.**

**Hepatocytes from AcSRSF1 HKO display widespread transcriptome defects.** **(A)** Histograms showing distribution of ΔPSI values of differentially spliced exons in acSRSF1 HKO with respect to controls from RNA-seq (n = 3 replicates per group). Distributions are separated by type of alternative exon event. Dotted vertical lines indicated median of distribution. **(B)** Volcano plot showing changes in mRNA abundance in acSRSF1 HKO. **(C)** Venn diagrams showing overlap of genes with differential expression and splicing in acSRSF1 HKO. **(D)** Breakdown of the number of SRSF1 eCLIP binding peaks located at the indicated pre-mRNA region. **(E)** Gene ontology analysis of the genes with SRSF1 binding peaks. **(F)** Top two enriched motifs for all SRSF1 eCLIP binding peak regions and for peaks associated to regulated gene sets. **(G)** Pie chart showing the percentage of SRSF1 binding peaks found to overlap with differentially expressed or spliced genes in acSRSF1 HKO. Bar plot on right shows the inverse relationship. **(H)** Cumulative peak density plot of SRSF1 binding peaks on differentially spliced cassette exons (green line) found in acSRSF1 HKO, native cassette exons (red), and constitutive exons (black) as background. The background set was sampled 1,000 times to determine the 95% confidence interval of peak density at each nucleotide position. Native cassette exon set consist of exons found to be alternative in both control and acSRSF1 HKO but did not change significantly ($0.05 < PSI < 0.95$ and $|\Delta PSI| \leq 0.05$) upon SRSF1 knockout.

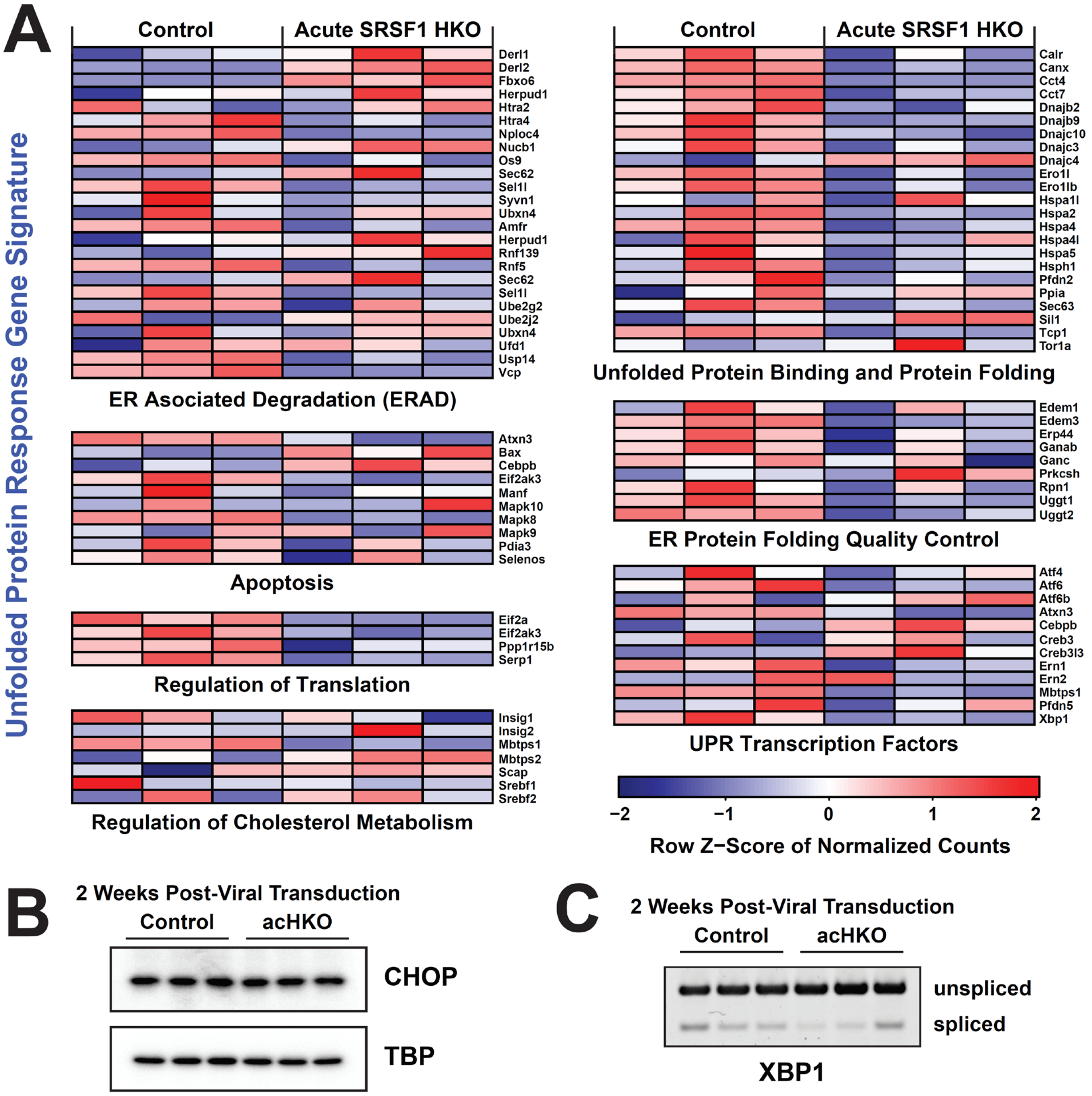


**Fig S6: Supplementary Figure S6 related to Figure 7.**

**Knockout of SRSF1 does not trigger the Unfolded Protein Response.** **(A)** Heatmap showing row normalized counts for genes involved in the Unfolded Protein Response, or UPR. Each column represents a single replicate. **(B)** Western blot showing CHOP levels in controls and acSRSF1 HKO with TBP serving as loading control. No significant difference in measured relative abundance. **(C)** XBP1 RT-PCR products showing relative levels of unspliced and spliced transcripts. No significant shift in splicing measured between control and acSRSF1 HKO.

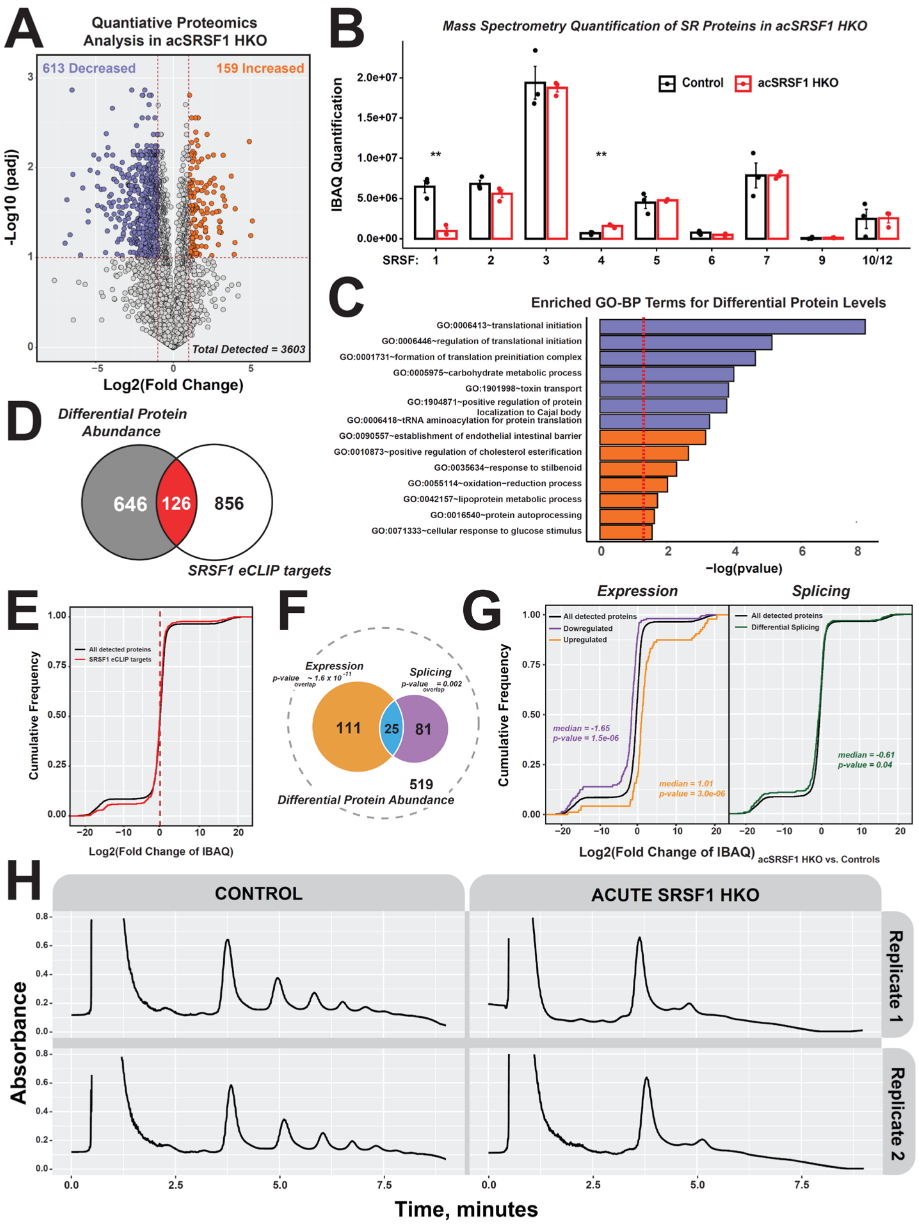


**Fig S7: Supplementary Figure S7 related to Figure 8.**

**Disruption of the hepatic proteome in acSRSF1 HKO.** **(A)** Plot depicting significant changes in relative protein abundances ($Log_2$|IBAQ ratio| >1, adjusted p-value < 0.1) estimated by quantitative mass-spectrometry based proteomics of control and acSRSF1 HKO (n = 3 per group). **(B)** GO analysis of differentially abundant proteins ($Log_2$(Fold Change) > 1, FDR < 0.10) in acSRSF1 HKO. **(C)** Plot of IBAQ quantification of detected SR proteins. **(D)** Overlap between differentially abundant proteins in acSRSF1 HKO and genes with SRSF1 eCLIP binding peaks. **(E)** Cumulative plot of protein fold-changes of all detected proteins (black) and subset with SRSF1 binding on associated gene. **(F)** Overlap of differentially abundant proteins with genes changing in expression or splicing with calculated overlap p-values. **(G)** Cumulative plot of protein fold-change values for associated genes that are differentially expressed (purple and orange for down- and up-regulated, respectively) or spliced (green) in acSRSF1 HKO. **(H)** Polysome profiles in hepatocytes from control and acSRSF1 HKO at 2-weeks post viral induction (n = 2 per group).

| Knockdown in HepG2 | Differential Expression (DESeq2) | | | Differential Splicing (rMATS) | | |
|---|---|---|---|---|---|---|
| | Upregulated | Downregulated | Total | Included | Skipped | Total |
| **SRSF1** | 1591 | 2312 | 3903 | 1999 | 1367 | 3366 |
| **SRSF3** | 834 | 376 | 1210 | 1087 | 624 | 1711 |
| **SRSF5** | 867 | 547 | 1414 | 729 | 667 | 1396 |
| **SRSF7** | 844 | 546 | 1390 | 433 | 587 | 1020 |
| **SRSF9** | 1272 | 1111 | 2383 | 476 | 865 | 1341 |

**Table S2: Supplementary Table S2 related to Figure 9.**
**Number of genes with differential expression or exons with differential splicing after shRNA knockdown of SR genes in HepG2 from the ENCODE Project.** DESeq2 and rMATS analysis was performed on ENCODE Project RNA-seq datasets for the indicated SR protein knockdown by shRNA in HepG2 cells. Differential expression and splicing were determined using parameter cut-offs as described previously.

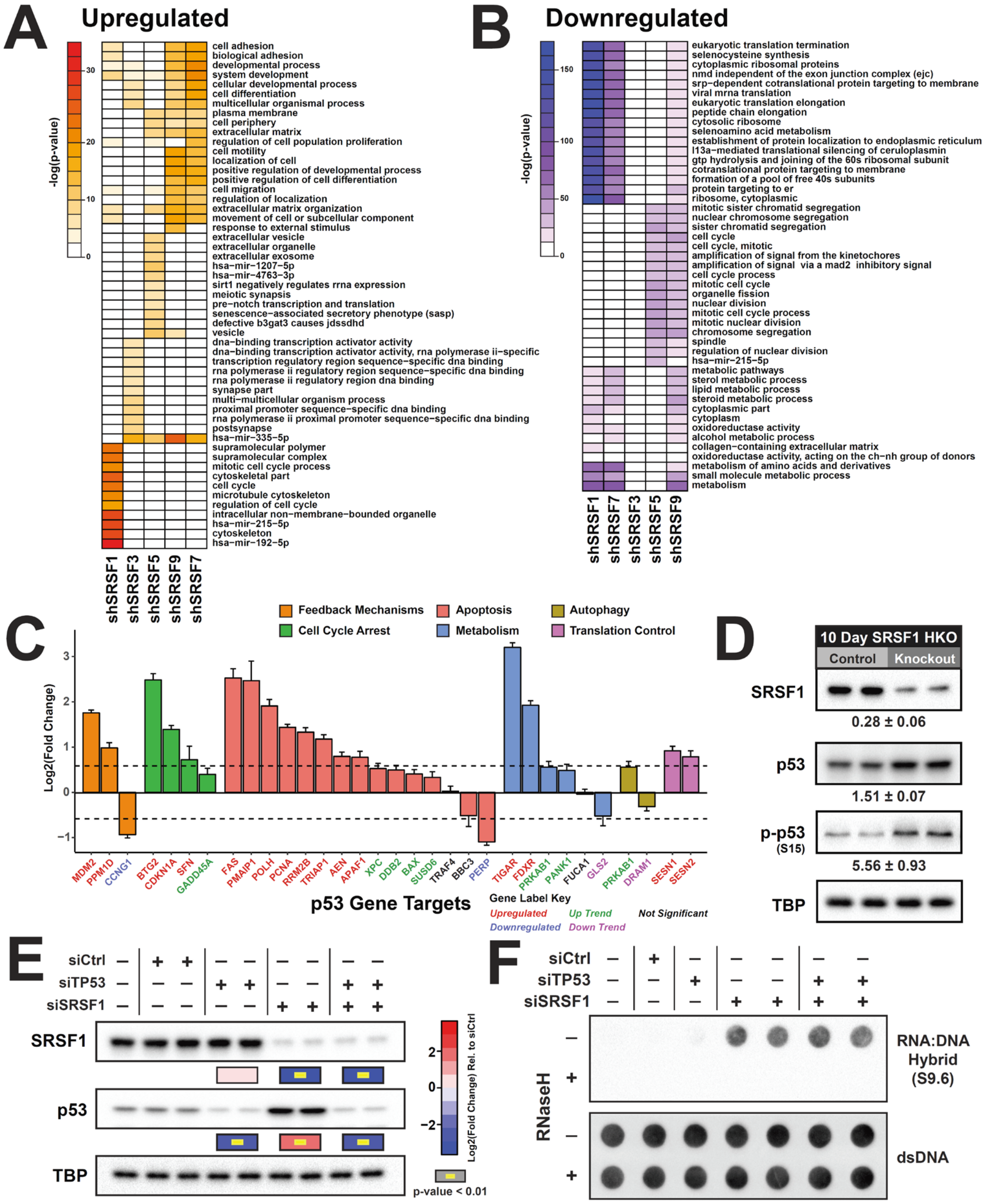


**Fig S8: Supplementary Figure S8 related to Figure 9.**

**Knockdown of SRSF1 in HepG2 triggers a p53 Response.** Heatmaps of gene ontology enrichment terms based on -log(p-value) for **(A)** upregulated and **(B)** downregulated genes in shRNA knockdown of SR proteins in HepG2 cells from the ENCODE Project. **(C)** Bar plot showing fold-changes in expression of various p53 target genes in HepG2 cells after SRSF1 knockdown (n = 2 per group). Color of x-axis gene labels depict direction of fold-change. **(D)** Western blot of 10-day control and SRSF1 HKO mice hepatocytes (n = 4 per group). Values under blot display mean ± SD of the abundance fold change in SRSF1 HKO relative to controls. **(E)** Western blots with quantifications represented below each blot in HepG2 cells with the indicated siRNA treatment (n=4 per group). **(F)** Representative R-loop dot blot on purified DNA from HepG2 cells with the specified siRNA knockdown (n = 4 per group). RNAse H treated samples serve as a negative control and a dsDNA dot blot was confirms equal loading.

| Antibody | Host Species | Type | Applications | Supplier Information, Catalog Number |
|---|---|---|---|---|
| anti-Mouse | Goat | Secondary, HRP | WB - 1:5000 | Bio-Rad, 1721011 |
| anti-Mouse, Dylight 488 | Goat | Secondary, Fluorescence | IF - 1:500 | Thermo Fisher Scientific, 35503 |
| anti-Mouse, Dylight 594 | Goat | Secondary, Fluorescence | IF - 1:500 | Thermo Fisher Scientific, 35511 |
| anti-Rabbit | Goat | Secondary, HRP | WB - 1:5000 | Thermo Fisher Scientific, 31460 |
| anti-Rabbit, Dylight 488 | Goat | Secondary, Fluorescence | IF - 1:500 | Thermo Fisher Scientific, 35553 |
| anti-Rabbit, Dylight 594 | Goat | Secondary, Fluorescence | IF - 1:500 | Thermo Fisher Scientific, 35561 |
| BAX | Rabbit | Primary | WB - 1:5000 | Abcam, ab32503 |
| Beta-actin | Rabbit | Primary | WB - 1:5000 | Cell Signaling Technology, 8457 |
| DNA-RNA Hybrid, Clone S9.6 | Mouse | Primary | DB - 1:5000 | Abcam, ab256361 |
| ds DNA | Mouse | Primary | DB - 1:5000 | Abcam, ab27156 |
| eIF2α | Rabbit | Primary | WB - 1:5000 | Cell Signaling Technology, 9722 |
| Hnf4a | Mouse | Primary | IF - 1:500 | Abcam, ab41898 |
| Hnf4a | Rabbit | Primary | IF - 1:500 | Cell Signaling Technology, 3113 |
| Ki67 | Mouse | Primary | IHC - 1:250 | BD Biosciences, 550609 |
| p53 | Rabbit | Primary | WB - 1:5000 | Abcam, ab131442 |
| phospho-eIF2α (S51) | Rabbit | Primary | WB - 1:5000 | Cell Signaling Technology, 9721 |
| phospho-gamma-H2A.X (S139) | Rabbit | Primary | IF - 1:500 | Abcam, ab11174 |
| phospho-p53 (S15) | Rabbit | Primary | WB - 1:5000 | Abcam, ab1431 |
| RIP | Rabbit | Primary | WB - 1:5000 | Abcam, ab106393 |
| SRSF1 | Rabbit | Primary | IF - 1:500, WB - 1:5000 | Abcam, ab129108 |
| SRSF1 (103) | Mouse | Primary | IF - 1:500, WB - 1:5000 | Invitrogen, 32-4600 |
| TBP | Mouse | Primary | WB - 1:10000 | Thermo Fisher Scientific, MA5-14739 |

**Table S3: Supplementary Table S3**

**Antibody dilutions and supplier information.** WB = western blot, IHC = immunohistochemistry, IF = immunofluorescence, DB = dot blot.

| Gene | Application | Forward Primer (5' → 3') | Reverse Primer (5' → 3') | Product Size |
|---|---|---|---|---|
| 36b4 | qPCR | AGATGCAGCAGATCCGCAT | GTTCTTGCCCATCAGCACC | 59 |
| Cre | qPCR | GCATTTCTGGGGATTGCTTA | ATTCTCCCACCGTCAGTACG | 95 |
| Xbp1 | Splicing | ACACGCTTGGGAATGGACAC | CCATGGGAAGATGTTCTGGG | 145, 171 |

**Table S4: Supplementary Table S4**
**Primer Sequences for qPCR and Splice Assay.**

| Experiment Target | Experiment Accession | Library-type | Utilized File Formats |
|---|---|---|---|
| SRSF7-human | ENCSR017PRS | paired-end | fastq, bam, bigWig |
| SRSF1-human | ENCSR094KBY | paired-end | fastq, bam, bigWig |
| SRSF9-human | ENCSR597XHH | paired-end | fastq, bam, bigWig |
| SRSF3-human | ENCSR376FGR | paired-end | fastq, bam, bigWig |
| SRSF5-human | ENCSR447UCG | paired-end | fastq, bam, bigWig |
| SRSF5-human | ENCSR781YNI | paired-end | fastq, bam, bigWig |
| Non-specific target control-human | ENCSR603TCV | paired-end | fastq, bam, bigWig |
| Non-specific target control-human | ENCSR264TUE | paired-end | fastq, bam, bigWig |
| Non-specific target control-human | ENCSR042QTH | paired-end | fastq, bam, bigWig |

**Table S5: Supplementary Table S5**
**SR Protein Knockdown in HepG2 ENCODE RNA-seq details.**

| GEO Accession Number | Model | Genotype | Treatment | Timepoint | Replicate | Total Number of Reads |
|---|---|---|---|---|---|---|
| GSM4412193 | SRSF1 HKO | AlbCre +/- | NA | 10 days | 1 | 101964963 |
| GSM4412194 | SRSF1 HKO | AlbCre +/- | NA | 10 days | 2 | 110546221 |
| GSM4412195 | SRSF1 HKO | SRSF1 fl/fl; AlbCre +/- | NA | 10 days | 1 | 116181095 |
| GSM4412196 | SRSF1 HKO | SRSF1 fl/fl; AlbCre +/- | NA | 10 days | 2 | 117337934 |
| GSM4412197 | SRSF1 HKO | AlbCre +/- | NA | 5 weeks | 1 | 117680056 |
| GSM4412198 | SRSF1 HKO | AlbCre +/- | NA | 5 weeks | 2 | 116619589 |
| GSM4412199 | SRSF1 HKO | SRSF1 fl/fl; AlbCre +/- | NA | 5 weeks | 1 | 115650894 |
| GSM4412200 | SRSF1 HKO | SRSF1 fl/fl; AlbCre +/- | NA | 5 weeks | 2 | 117418842 |
| GSM4412201 | acSRSF1 HKO | SRSF1 fl/fl | AAV8-TBG-GFP | 2 weeks | 1 | 127573517 |
| GSM4412202 | acSRSF1 HKO | SRSF1 fl/fl | AAV8-TBG-GFP | 2 weeks | 2 | 176265351 |
| GSM4412203 | acSRSF1 HKO | SRSF1 fl/fl | AAV8-TBG-GFP | 2 weeks | 3 | 139425979 |
| GSM4412204 | acSRSF1 HKO | SRSF1 fl/fl | AAV8-TBG-iCRE | 2 weeks | 1 | 119934148 |
| GSM4412205 | acSRSF1 HKO | SRSF1 fl/fl | AAV8-TBG-iCRE | 2 weeks | 2 | 139028073 |
| GSM4412206 | acSRSF1 HKO | SRSF1 fl/fl | AAV8-TBG-iCRE | 2 weeks | 3 | 115531464 |

**Table S6: Supplementary Table S6**
**SRSF1 HKO Model RNA-seq Sample Information.**
